# A rapid CRISPR competitive assay for *in vitro* and *in vivo* discovery of potential drug targets affecting the hematopoietic system

**DOI:** 10.1101/2021.03.08.434360

**Authors:** Yunbing Shen, Long Jiang, Vaishnavi Srinivasan Iyer, Bruno Raposo, Sanjay V. Boddul, Zsolt Kasza, Fredrik Wermeling

## Abstract

CRISPR/Cas9 can be used as an experimental tool to inactivate genes in cells. However, a CRISPR-targeted cell population will not show a uniform genotype of the targeted gene. Instead, a mix of genotypes is generated - from wild type to different forms of insertions and deletions. Such mixed genotypes complicate analyzing the role of the targeted gene in the studied cell population. Here, we present a rapid experimental approach to functionally analyze a CRISPR-targeted cell population that does not involve generating clonal cell lines. As a simple readout, we leverage the CRISPR-induced genetic heterogeneity and use sequencing to identify how different genotypes are enriched or depleted related to the studied cellular behavior or phenotype. The approach uses standard PCR, Sanger sequencing, and a simple sequence deconvoluting software, enabling laboratories without specific in-depth knowledge to also perform these experiments. As proof of principle, we present examples studying the role of different genes for various aspects related to hematopoietic cells (T cell development *in vivo* and activation *in vitro*, macrophage phagocytosis, and a leukemia-like phenotype induced by overexpressing a proto-oncogene). In conclusion, we present a rapid experimental approach to identify potential drug targets related to mature immune cells, as well as normal and malignant hematopoiesis.

**Highlights:**

‐ CRISPR generates genetic heterogeneity at the targeted site.
‐ Genetic heterogeneity complicates identifying the role of a targeted gene.
‐ Heterogeneity can be quantified by Sanger sequencing with sufficient sensitivity.
‐ Enrichment of specific genotypes can be used to identify roles for targeted genes.
‐ Competitive experiments show the potential of genotype enrichment as a discovery tool.

**Graphical representation:** 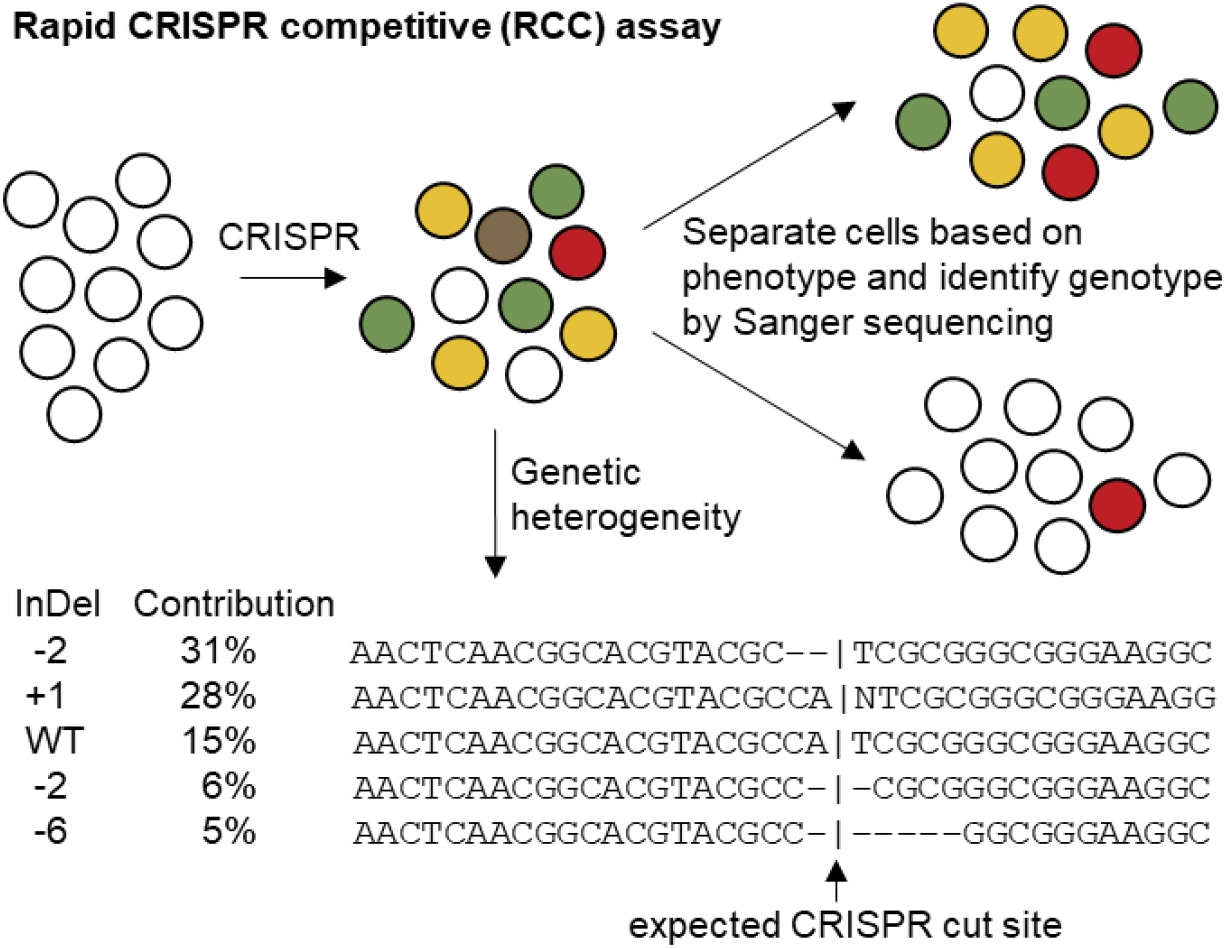

## 1. Introduction

### 1.1 Clustered Regularly Interspaced Short Palindromic Repeats (CRISPR)

CRISPR has been developed from its natural prokaryotic origins into a set of molecular biology tools that can be used to modify genes in eukaryotic cells [1-4]. In its most simple form, a CRISPR experiment involves delivering a single guide RNA (sgRNA), with specificity for the gene of interest, and the endonuclease Cas9 into the nucleus of the studied cell [5]. Due to its simplicity, CRISPR is playing an increasingly important role in generating cell lines and animal models with specific genetic modifications. Experiments comparing pairs of cell lines or animals that differ at one specific genetic region, for example being wild type (WT) and knockout (KO) for a gene of interest, is a powerful approach extensively used to identify the role of the gene for a studied phenotype. However, the mutation spectrum generated in a CRISPR-targeted cell population is not uniform. Instead, both unmodified (WT), as well as insertions and deletions (InDels) of different sizes, are typically generated when the Cas9 induced DNA damage is repaired by the error-prone non-homologous end-joining pathway [6, 7]. The genetic heterogeneity makes it difficult to directly analyze the role of a targeted gene, and researchers often generate clonal lines with defined mutations from the modified cell population. It is, however, not feasible to generate extensive clonal lines from many cell types, including most primary cell populations. Several approaches have been developed to evaluate the genetic heterogeneity in a CRISPR-targeted cell population. These include next-generation sequencing (NGS) platforms [8-10], approaches based on fragment length analysis (FLA) of PCR amplicons, like IDAA (Indel Detection by Amplicon Analysis) [11, 12], as well as analysis tools like ICE (Inference of CRISPR Edits) [13], and TIDE (Tracking of Indels by DEcomposition) [14] that deconvolute Sanger sequencing data into the frequency of different genotypes found in a sample.

### 1.2 The hematopoietic system

Hematopoiesis is the essential process where mature immune cells, platelets, and erythrocytes are formed from hematopoietic stem cells (HSCs) located in specific adult bone marrow (BM) niches of higher vertebrates [15-18]. This concept was formally proven in the 1950s by experiments and clinical treatment showing that the transplantation of BM cells into an irradiated host results in the formation of mature cells stemming from the donor HSCs [19, 20]. Due to its feasibility, BM transplantations have been extensively used in experimental immunological research, for example, to compare the response of immune cells with different genotypes *in vivo*. As such, BM cells from mice with different genotypes (for example WT and KO for a gene of interest) can be combined and transplanted into an irradiated recipient mouse, generating a “mixed BM chimeric” mouse. By using different congenic markers, like CD45.1 and CD45.2 [21, 22], to track cells from the different BM donors, cells with different genotypes in the recipient mouse can be separated by flow cytometry and the role of the targeted gene identified for a studied phenotype.

Of additional importance, malignancies at different developmental stages of the hematopoietic lineage, including leukemia, represent the major cancer types seen in children and adolescence [23], as well as constituting a significant amount of all cancers observed in adults [24].

## 2. Material and methods

### 2.1 Mice

8-to 12-week-old, sex- and age-matched mice were used in experiments. All mice were housed in specific pathogen-free conditions with a 12/12-hour light/dark cycle and fed standard chow diet ad libitum. All animal experiments were approved by the local ethical committee at Karolinska Institute, Sweden. The following mouse strains from Jackson Laboratory were used: C57BL/6 Cas9+ GFP+ (stock no. 026179, CD45.2+), and C57BL/6 CD45.1 (stock no. 002014). C57BL/6 Cas9+ GFP+ mice and CD45.1 mice were crossed, detecting GFP and CD45.1 by flow cytometry, to generate homozygous Cas9+ GFP+ CD45.1+ mice (Cas9.1).

Bone marrow transplantation was performed by i.v. injection of ∼10^6^ bone marrow cells into recipient mice irradiated with 900 rad of *γ*-irradiation 12-24 hours earlier. The bone marrow cells were typically electroporated with a sgRNA just before being injected into the recipient mice. To evaluate the mutations of the BM cells, a fraction of the electroporated cells were kept in culture, to allow for the CRISPR event to occur, and sequenced two days later.

### 2.2 sgRNA and primer design

The Green Listed software (http://greenlisted.cmm.ki.se) [25, 26] utilizing the Brie reference library, typically selecting the sgRNA with the highest on-target activity [27], or https://design.synthego.com/#/ were used to design sgRNAs. sgRNAs with stabilizing 2’-O-methyl and phosphorothioate linkages were ordered from Sigma-Aldrich or Synthego. The geneMANIA plugin for Cytoscape [28] was used to identify potential interaction partners of HOXB8, as discussed in [26]. Primers were designed using Primer-BLAST (https://www.ncbi.nlm.nih.gov/tools/primer-blast/), aiming for a 400-800 bp amplicon with the sgRNA binding site in the middle. The used sgRNA and primer sequences are listed in Supplementary tables 1 and 2.

### 2.3 Isolating, CRISPR modifying, and differentiating bone marrow (BM) cells

BM cells were collected by flushing femurs and tibias with PBS. Lineage negative cells (Lin-) were obtained by depleting lineage positive cells (Lin+) from the BM cells using MACS buffer (Miltenyi Biotec, #130-091-221), Lineage Cell Detection Cocktail-Biotin (Miltenyi Biotec, #130-092-613, 1:100), Anti-Biotin MicroBeads (Miltenyi Biotec, #130-090-485), and LS column (Miltenyi Biotec, #130-042-401), according to the protocol suggested by the manufacturer. Lin-cells were culture in complete RPMI medium (cRPMI) containing 20 ng/ml of SCF (PeproTech, #250-03), TPO (PeproTech, #315-14), IL-3 (PeproTech, #213-13), and IL-6 (PeproTech, #216-16) for two days. cRPMI: RPMI-1640 (Sigma-Aldrich #R0883) with 10% heat-inactivated fetal bovine serum and 1% penicillin-streptomycin-glutamine (100X, Gibco, #10378016). Cells were cultured at 37 °C in a humidified incubator with 5% CO_2_ and handled in laminar flow hoods using standard sterile techniques.

The Neon Transfection System (Invitrogen, #MPK5000) was used for electroporation-based delivery of CRISPR components, following the manufacturer’s instructions initially using the suggested program testing 24 different conditions. Electroporation condition #5 (Pulse voltage: 1700 V, Pulse width: 20 ms, Pulse number: 1) was used unless otherwise specified. For BM cells, typically 50-100 pmol sgRNA was delivered into 2*10^5^ cells per electroporation using the Neon 10 µL Kit (Invitrogen, #MPK1096) or 500-1000 pmol of sgRNA delivered into 2*10^6^ cells per electroporation using the Neon 100 µL Kit (Invitrogen, MPK10096). Electroporated Lin-cells were kept in culture for two days in cRPMI with cytokines before sequencing to allow for the CRISPR events to occur. Trp53 siRNA was typically delivered in the same reaction as the sgRNAs. Trp53 ON-TARGETplus mouse siRNA SMARTPool was ordered from Horizon Discovery (#L-040642-00-0005). 100 pmol of siRNA was delivered into 2*10^5^ Lin-cells per electroporation experiment.

To differentiate the BM cells in vitro, electroporated Lin-cells were switched to indicated cytokines directly after electroporation; for macrophages, cRPMI with 100 ng/ml of M-CSF (PeproTech, #315-02) and cultured for 7 days, exchanging half the medium every 2-3 days; for dendritic cells, cRPMI with 100 ng/ml of Flt3L (Biolegend, #550706) and cultured for 9 days, with one 1:2 split after 4-5 days.

The macrophage phagocytosis assay was performed using a kit (Cayman Chemical, #600540) as suggested by the manufacturer. Briefly, differentiated macrophages were incubated with the Latex Beads-Rabbit IgG-PE complex (1:250) in a 6 well plate with 3 ml of cRPMI for 3 hours at 37 °C. Cells were then washed gently and collected for further analysis.

### 2.4 Generating and culturing Hoxb8 BM cells

The Hoxb8 cells were generated by transducing bone marrow cells of C57BL/6 Cas9+ GFP+ mice with an estrogen-inducible retroviral construct expressing HOXB8 (ER-Hoxb8, a kind gift from Mark P. Kamps, University of California, San Diego) as described [29, 30]. Transduced BM cells were cultured in 1 µM β-estradiol (BE, Sigma-Aldrich, #E2758) and 25 nM mouse SCF (PeproTech, #250-03) for several weeks with HOXB8 expression turned on to establish a cell line-like population. To inactive the HOXB8 activity, BE was withdrawn from the media for 3 days. The Hoxb8 cells were CRISPR modified in the same way as BM cells (described in 2.2).

### 2.5 Culturing and modifying peripheral blood mononuclear cells (PBMC) and Jurkat cells

PBMCs were derived from buffy coats from consenting healthy donors (Karolinska Hospital Blood Bank). PBMCs were isolated using Ficoll-Paque Plus (GE Lifesciences, #17144002) according to the manufacturer’s recommended protocol. PBMCs were cultured in CTS OpTmizer (Gibco, #A1048501) with 10% heat-inactivated fetal bovine serum, 1% penicillin-streptomycin-glutamine and 25 units/mL of IL-2 (Peprotech, #200-02), exchanging half the medium every 2-3 days. To expand the T cell population, PBMCs were stimulated with CD3/28 beads (Milteny Biotech, #130-091-441), re-stimulated every 7-10 days, and analyzed by flow cytometry to confirm the percentage of T cells in the culture. When used for sgRNA electroporation, culture was more than 90% T cells (TCR-α/β positive cells by flow cytometry).

The Jurkat-NFAT-GFP cell line was generated by transducing Jurkat cells (ATCC, TIB-152) with the pSIRV-NFAT-eGFP plasmid (Addgene, #118031, a gift from Peter Steinberger [31]) as described in Boddul et al. [32], with the modification that Ecotropic Receptor Booster (Takara, #631471) was added to the cells as suggested by the manufacturer. The cells were maintained in cRPMI.

For both the Jurkat and PBMCs, the Neon electroporation condition #24 (Pulse voltage: 1600 V, Pulse width: 10 ms, Pulse number: 3) was used. 60 pmol of sgRNA was complexed with 10 pmol of Cas9 protein (Sigma-Aldrich, #CAS9PROT) and electroporated into 0.5*10^5^ Jurkat cells per reaction, and 100 pmol of sgRNA was complexed with 16 pmol of Cas9 protein and electroporated into 2*10^5^ PBMCs per reaction using the Neon Transfection System 10µL Kit.

WT, electroporated (empty) control, and T cell receptor alpha chain constant (TRAC) sgRNA electroporated PBMCs or Jurkat cells were cultured for at least seven days before the experiment. The cells were stimulated with 100 nM PMA/Ionomycin (Sigma-Aldrich, #P8139 and #I3909) or CD3/28 beads (Milteny Biotech, #130-091-441) in a 1:1 bead to cell ratio for 18 hours and the beads were removed with the MACSiMAG Separator (Milteny Biotech, 130- 092-168), before analysis by flow cytometry.

### 2.6 Flow cytometry analysis and sorting

Single-cell suspensions were stained for 30 min, washed and sorted using Sony SH800S, or acquired using BD LSRFortessa, BD FACSVerse, BD Accuri, or Cytek Aurora. Generated FCS files were analyzed by FlowJo version 10 (FlowJo, LLC).

Lin-BM cells were stained with Sca1–PE/Cy7 (Biolegend, #108113), c-Kit–APC (BD Biosciences, #561074), Lin–biotin (Lineage Cell Detection Cocktail-Biotin, Miltenyi Biotec, #130-092-613), Streptavidin–PE (BD Biosciences, #554061) and LIVE/DEAD Fixable Aqua Dead Cell Stain Kit (Invitrogen, #L34957).

B cells and T cells were sorted from Zap70 iCR mice spleen stained with CD45.1–FITC (BD Biosciences, #561871), CD45.2–BV785 (Biolegend, #109839), TCRb–BV711 (Biolegend, #109243), B220–PE (Invitrogen, #12-0452-82) and DAPI (Sigma-Aldrich, #D9542, 0.1 µg/ml).

Macrophages were stained with CD11b–PerCP/Cy5.5 (Biolegend, #101228), F4/80–PE (Biolegend, #123110), and DAPI.

Dendritic cells were stained with CD11c-PE-Cy7 (Biolegend, #117318), I-A/I-E-AlexaFluor 647 (Biolegend, #107617), CD80-APC (Biolegend, #104713), CD86-FITC (Biolegend, #105005), CD274-PE (Biolegend #124307) and LIVE/DEAD Fixable Near-IR dead cell stain kit (ThermoFisher Scientific #L10119). After 15 min of staining at room temperature, cells were washed and analyzed by flow cytometry machine Cytek Arora.

Hoxb8 cells were stained with biotin anti-mouse Lineage Panel (BioLegend #133307), Streptavidin-BV421 (BD Biosciences, #563259), LIVE/DEAD Fixable Aqua Dead Cell Stain Kit (Invitrogen, #L34957).

To confirm TCRα knockout efficiency on Jurkat cells and PBMCs, cells were stained with TCR α/β–APC antibody (Biolegend, #306717). Stimulated Jurkat cells and PBMCs were stained with TCR α/β–APC antibody (Biolegend, #306717) CD69-PE (Biolegend, #310905), and LIVE/DEAD Fixable Aqua Dead Cell Stain Kit (Invitrogen, #L34957).

### 2.7 Sanger sequencing, Inference of CRISPR Edits (ICE) analysis, and Indel Detection by Amplicon Analysis (IDAA)

At least 10,000 sorted cells or 10 μL of whole blood sample were collected for genomic DNA extraction using the DNeasy Blood & Tissue Kit (Qiagen, #69504). 3 μL of genomic DNA was used as template to amplify the sgRNA target region, using a standard PCR program. Amplicons were purified directly from PCR reaction mix by using DNA Clean & Concentrator Kits (Zymo Research, #D4013) or recovered from agarose gel by using Zymoclean Gel DNA Recovery Kit (Zymo Research, #D4007). The PCR products were quantified by Nanodrop and sequenced by Eurofins Genomics. The Sanger sequencing data was subsequently analyzed by ICE (Synthego, https://ice.synthego.com). For the IDAA fragment length analysis, genomic DNA samples were sent to COBO Technologies (https://cobotechnologies.com/).

### 2.8 Statistics

Statistical tests were performed as indicated in the respective figure legend using GraphPad Prism 8.

## 3. Theory/calculation

Sanger sequencing can be used as a simple readout to identify the role of CRISPR-targeted genes in complex cellular behaviors.

## 4. Results

### 4.1 Lineage negative (Lin-) bone marrow (BM) cells can readily be modified by CRISPR and evaluated by sequencing

To enable studying the role of different genes in the hematopoietic system, we first set out to optimize modifying HSCs with CRISPR. To this end, we isolated BM cells from Cas9+ GFP+ mice on the C57BL/6 background [33]. HSCs were enriched by lineage (Lin) depletion (to eliminate mature Lin+ cells) and electroporated with a GFP targeting sgRNA. The extent of GFP inaction (KO) was analyzed by flow cytometry (**Fig. 1A**). Screening different electroporation programs, we identified a set of parameters that gave a good KO efficiency without a substantial effect on cell survival (**Fig. 1B-C**). We selected condition #5 (pulse voltage: 1700 V, pulse width: 20 ms, pulse numbers: 1) and further tested how the concentration of sgRNA affected the KO efficiency, identifying that doses >50 pmol gave a high and uniform KO efficiency (**Fig. 1D**). Next, we tested additional parameters affecting the KO efficiency, including different storage conditions of the sgRNA, as well the inclusion of Trp53 siRNAs, since transient p53 inhibition has been shown to increase the KO efficiency in CRISPR experiments [34, 35] while protecting the function of HSCs [36]. We found that using freshly prepared sgRNAs and Trp53 siRNA increased the KO efficiency of the targeted gene (**Fig. 1E**). GFP, as well as surface markers that can be readily stained with antibodies, are easily followed by flow cytometry. However, the genotype of most genes is not easily evaluated by flow cytometry. As an alternative readout, we hypothesized that we instead could use standard Sanger sequencing to quantify the CRISPR-induced genotype. Using the ICE software [13] to analyze Sanger sequencing data, we identified that the GFP targeting sgRNA used generated a diverse genotype in the Lin-cells, with a dominant +1 insertion next to the expected cut site (**Fig. 1F**). To compare the two methods assessing CRISPR-efficiency of gene KO, we generated a dilution curve of cells with different levels of GFP KO by diluting sgRNA electroporated Lin-BM cells with different proportions of non-electroporated Lin-BM cells. Sequencing of the mutation frequency was then compared to the KO phenotype identified by flow cytometry in the same cells. We found a good correlation between the two readouts (R^2^ = 0.87, p = 0.0007), although the sensitivity of the sequencing readout was decreased when mutations were found at a low frequency, something that can be expected by the nature of the sequence deconvolution (**Fig. 1G**). As an alternative, we used the IDAA fragment length analysis approach and found a very strong correlation to the flow cytometry readout (R^2^ = 0.99, p < 0.0001) (**Fig. 1H**). Sequencing and IDAA can thus both be used as readouts to quantify the mutation frequency when flow cytometry is not a feasible readout.

**Figure 1.**
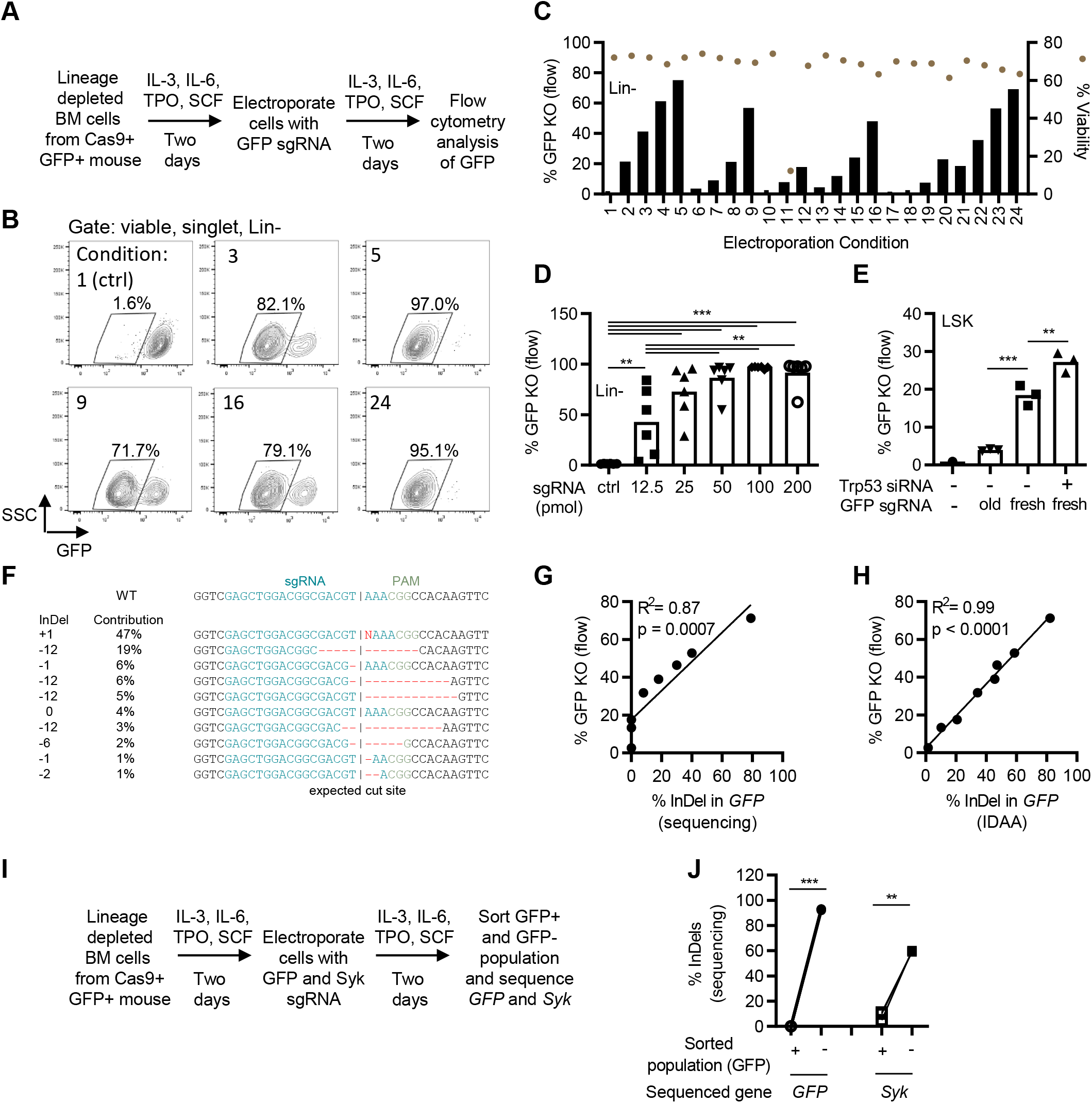
Lineage negative (Lin-) bone marrow (BM) cells can readily be modified by CRISPR and evaluated by sequencing. (**A**) Model describing the experimental setup where Lin-BM cells were cultured in a cytokine cocktail, electroporated with a GFP targeting sgRNA, and analyzed by flow cytometry. (**B-C**) flow cytometry analysis of GFP expression in Lin-BM cells two days after electroporation with a GFP sgRNA comparing different electroporation programs. Condition 5 (pulse voltage: 1700 V, pulse width: 20 ms, pulse numbers: 1) was selected for further experiments. B and C represent two different experiments with slightly different efficiency. (**D**) Flow cytometry analysis of GFP KO cell % two days after electroporation with different amounts of the GFP sgRNA. Gate: viable, single, Lin-cells. (**E**) Flow cytometry analysis of GFP KO cell % two days after electroporation with a sgRNA solution that had been stored in -20°C for several weeks (old), or a freshly dissolved sgRNA (fresh) in the presence of a Trp53 siRNA. Gate: viable, single, Lin-, Sca-1+, c-Kit+ (LSK) cells. (**F**) Analysis of the sgRNA targeted GFP region showing the generated mutation spectrum. Data generated by the ICE software based on Sanger sequencing of PCR product. (**G**) Comparison of the identified % insertion and deletions (InDels) in GFP using sequencing and the % GFP KO cells by flow cytometry. (**H**) Comparison of the identified % insertion and deletions (indels) in GFP using the InDel Detection by Amplicon Analysis (IDAA) assay and the % GFP KO cells by flow cytometry. (I) Model describing simultaneously targeting *GFP* and *Syk*. (**J**) Quantification of % InDels in *GFP* and *Syk* in cells sorted based on GFP expression, Data shown as individual data points (B-C, G-H, J, n=3), mean and individual data points (D, n=6; E, n=3). ** = p < 0.01, *** = p < 0.001 by one-way ANOVA and Turkey’s post-test (D-E), simple linear regression (G-H), or paired T-test (J).

We next hypothesized that the mutation frequency of one targeted gene (*X*), could predict the mutation frequency of another gene (*Y*) in a cell population simultaneously electroporated with two sgRNAs (targeting *X* and *Y*). Such an approach is based on the idea that if a cell inactivates one gene, it has a high chance of also successfully inactivate another co-targeted gene. This type of approach could be used as a strategy to enrich for cells with the intended mutation, similar to what has been described by co-targeting *DTR* [37], or *HPRT* [38]. To this end, we electroporated the Lin-BM cells with a combination of GFP and Syk sgRNAs, sorted the GFP positive (+) and GFP negative (-) cells after two days, and sequenced the targeted *GFP* and *Syk* loci in the sorted cells (**Fig. 1I**). In line with the hypothesis, cells that failed to inactivate GFP (GFP+ cells), had no detectable mutations in *GFP*, and only minimal in *Syk* (**Fig. 1J**). By now, we concluded that: (*i*) we had established an optimized system for modifying genes in Lin-BM cells by CRISPR, (*ii*) a simple Sanger sequencing readout could be used to quantify the mutation frequency in a cell population, albeit not when the mutations are found at a low frequency, and (*iii*) co-targeting *GFP* and a second gene of interest followed by sorting cells with inactivated GFP constitutes a strategy to enrich for mutations in the gene of interest.

### 4.2 Generating immuno-CRISPR (iCR) mice and evaluating the CRISPR-mediated modifications by sequencing

Next, we applied the optimized protocol to generate *in vivo* models with the modified Lin - BM cells. For this purpose, Lin-BM cells from Cas9+ mice were cultured and electroporated with a Zap70 targeting sgRNA, followed by transplantation to irradiated recipients, generating what we refer to as immuno-CRISPR (iCR) mice (**Fig. 2A**). ZAP70 is a component of the T cell receptor signaling pathway essential for mature T cell development [39, 40], but with a redundant role for B cell development [41]. In line with the literature related to *Zap70* deficiency, we saw a diminished T cell population in the spleen of Zap70 iCR mice (**Fig. 2B-C**). Furthermore, after sequencing *Zap70* in sorted B and T cells from the Zap70 iCR mice, we observed that the B cells showed a high mutation frequency, in concordance with that ZAP70 is not important for B cell development. In contrast, none of the sorted T cells had any detectable *Zap70* mutations (**Fig. 2D**).

**Figure 2.**
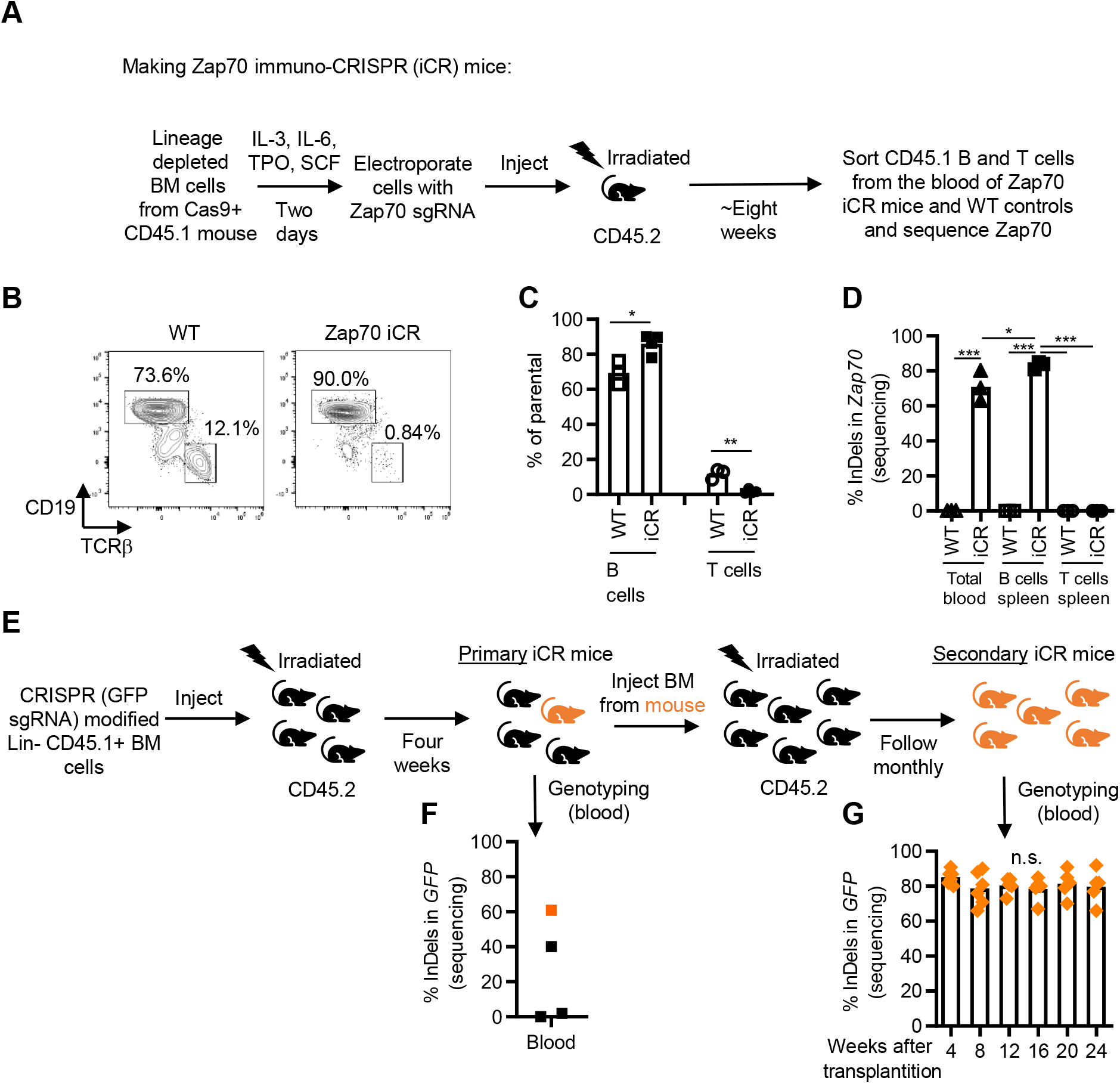
Generating immuno-CRISPR (iCR) mice and evaluating the CRISPR-mediated modifications by sequencing. (**A**) Model describing the experimental setup where CD45.1+ Lin-BM cells were modified by CRISPR targeting *Zap70* and grafted into irradiated CD45.2+ recipients. (**B**) Flow cytometry analysis of cells in the blood of Zap70 iCR mice and WT control mice eight weeks post transplantation. Cells gated on viable, CD45.1+, single lymphocytes. (**C**) Quantification of B and T cells in the blood of WT and Zap70 iCR mice in (B). (**D**) Analysis of the level of mutations in the sgRNA targeted *Zap70* region in total cells from the blood, as well as in B and T cells sorted from the spleen of Zap70 iCR mice and WT control mice. (**E**) Model describing the experimental setup where a secondary transplantation was used to amplify the population of successfully modified mice. (**F**) Analysis of the level of mutations in the sgRNA targeted *GFP* region in blood cells of the GFP iCR mice four weeks after transplantation, in an experiment with low efficiency. One mouse showed good knockout efficiency (labeled in orange) and was used as BM donor for secondary transplantation. (**G**) Kinetics of the level of mutations of GFP in the secondary iCR mice. Data shown as mean and individual data points (C-D, n=3; F, n=4; G, n=5-6). n.s. = non-significant, * = p < 0.05, ** = p < 0.01 *** = p < 0.001 by unpaired T-test (C), and one-way ANOVA and Turkey’s post-test (D, H).

Occasionally, the mutation frequency achieved in CRISPR-targeted Lin-BM cells is low, and as a consequence iCR mice generated from these cells typically have a low frequency of mutations and a higher level of variability between recipient mice as exemplified in **Fig. 2E-F**. We have noted that performing secondary transplantations from a single successful iCR mouse in such a situation can expand the number of mice with the desired mutation (**Fig. 2G**). Importantly, this also gives an example of how an iCR mouse population could be expanded by secondary transplantation, something that is considerably faster than expanding a traditional colony of mice by breeding. We concluded that sgRNA electroporated Lin-HSCs can be grafted into irradiated recipient mice, resulting in the formation of mature immune cells carrying the intended mutation. Furthermore, the role of a targeted gene in the differentiation of mature immune cell s *in vivo* can be evaluated by sequencing, comparing the genotype of different cell populations.

### 4.3 *In vitro* differentiation of CRISPR-modified BM cells into macrophages and dendritic cells

In immunological research, immature BM cells are commonly differentiated *in vitro* into different mature myeloid immune cell populations by the addition of specific cytokines to the cell culture medium [42]. This setup allows for controlled experiments testing parameters in isolated, non-transformed, immune cells. We next investigated whether the CRISPR-modified Lin-BM cells could be differentiated *in vitro* to defined mature immune cell populations by culturing the cells in M-CSF (for macrophage differentiation), or F lt3L (for dendritic cell differentiation) (**Fig. 3A**). In the M-CSF culture, we observed a good differentiation of cells into the expected F4/80^high^ CD11b^high^ macrophage phenotype and observed a high degree of GFP KO efficiency (**Fig. 3B-C**). To assess whether the functionality of the macrophages was non-specifically affected by the CRISPR modification, we added PE-labeled IgG-coupled beads to assess phagocytosis by these cells (**Fig. 3D**). Importantly, we found no difference in the GFP genotype in the sorted PE^high^ and PE^low^ cells (**Fig. 3E**). A successful KO event in an irrelevant gene (GFP in this case) did thus not affected the cells in a non-specific way, something that could be considered related to how DNA damage affects cells.

**Figure 3.**
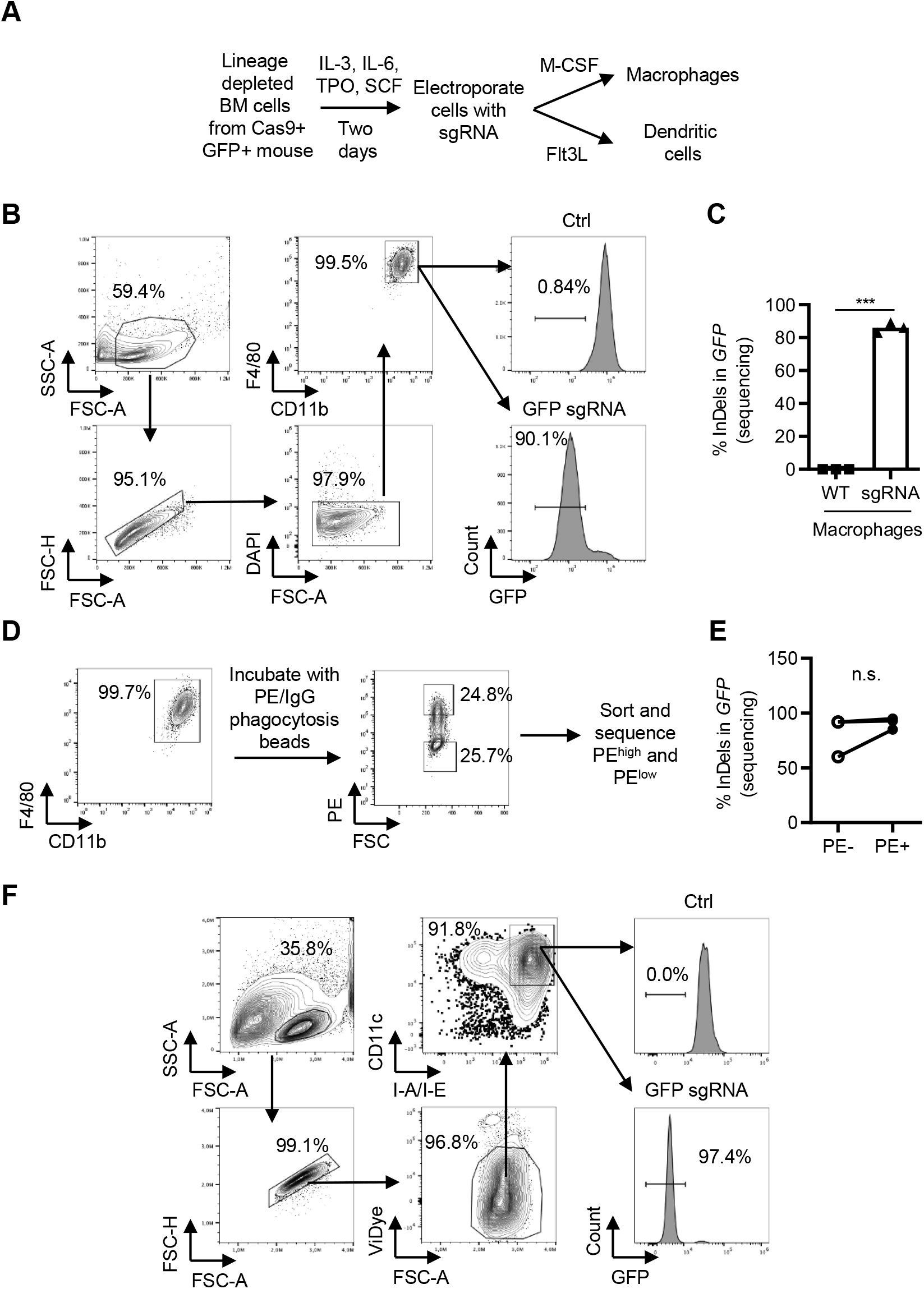
*In vitro* differentiation of CRISPR-modified BM cells into macrophages and dendritic cells. (**A**) Model describing the experimental setup. (**B**) Flow cytometry plots showing the gating strategy for cells differentiated for seven days in M-CSF. (**C**) *GFP* InDel frequency in sorted viable macrophages from M-CSF cultures. (**D**) Macrophages were incubated with PE/IgG phagocytosis beads and three hours later sorted for the level of binding to the beads into PE^high^ and PE^low^ populations. (**E**) The sorted populations were sequenced and the % of *GFP* InDels in the two populations was quantified. (**F**) Flow cytometry plots showing the gating strategy for cells differentiated for nine days in Flt3L. Data shown as representative flow cytometry plots (B, D, F), mean and individual data (C, n=3), and individual data (E, n=3). n.s. = non-significant, *** = p < 0.001 by unpaired T-test (C), and paired T-test (E).

Similarly, in the Flt3L-supplemented culture, we observed that the CRISPR-modified Lin-BM cells differentiated well into dendritic cells (CD11c^high^, MHC II^high^) and showed a good level of GFP KO efficiency (**Fig 3F**). In addition, instead of modifying precursor cells, that are subsequently differentiated to mature immune cell populations, the sgRNA can also be electroporated directly into mature immune cells, as exemplified with a human T cell line, and peripheral blood mononuclear cells (PBMC) in **Supplementary Fig. 1**. We concluded that (*i*) the CRISPR-modified Lin-BM cells could be successfully differentiated into macrophages and dendritic cells, and that (*ii*) competitive functional assays can be performed with the cells to identify how specific genes are affecting a studied behavior (as exemplified by phagocytosis), using a sequence-based readout.

### 4.4 Using the Rapid CRISPR Competitive Assay (RCC) to study transformation by the HoxB8 proto-oncogene

Lastly, we wanted to assess if our experimental setup could be used to study the role of different genes in relation to malignancies of the hematopoietic linage. To this end, we transduced the Cas9+ GFP+ BM cells with an inducible construct expressing the proto-oncogene *Hoxb8* [43] and electroporated them with different sgRNAs to identify the role of targeted genes for the HOXB8 transformed phenotype (**Fig. 4A**). When the activity of HOXB8 is induced, the BM cells are proliferating at an immature Lin-stage with a granulocyte-macrophage precursor (GMP) phenotype (**Fig. 4B**) [29]. As such the cells show behavioral (unlimited proliferation, block in differentiation) and phenotypic (immature, Lin-) features that overlap with acute leukemia cells as has been proposed [44, 45]. In contrast, when the HOXB8 activity is turned off, the HOXB8-induced proliferation and differentiation block are eliminated, and the cells differentiate into mature, Lin+, cells with a limited lifespan (**Fig. 4C**) [29, 46]. To test the experimental setup, we electroporated the HOXB8 transformed cells with a HoxB8 sgRNA and three days later observed that approximately 50% of the cells had acquired the Lin+ phenotype, expected when the activity of HOXB8 was turned off (**Fig. 4D**). We subsequently sorted the cells into Lin - and Lin+ and sequenced the *Hoxb8* locus to quantify the mutation spectrum. As expected, we found that the Lin+ population, which behaved as if the HOXB8 activity was turned off, showed a 100% mutation frequency in *Hoxb8* (**Fig. 4E**). To our surprise, we also noted that the Lin-population had a significant amount of mutations in the *Hoxb8* gene. Detailed analysis showed that these mutations almost exclusively corresponded to different insertion or deletion (InDels) with a multiplier of three nucleotides, in contrast to the Lin+ population where InDels consisted of a multiplier of one or two nucleotides (**Fig. 4E-F**). This is in line with the fact that an InDel with a multiplier of one or two nucleotides causes a frameshift, premature stop codons, and nonsense-mediated decay, essentially causing a KO of the gene in most cases [47, 48]. On the other end, InDels with a multiplier of three nucleotides will result in the insertion or deletion of amino acids (AA) in the translated protein, something that depending on the protein and the specific site where the change occurs, can inactivate the protein, or, as it appears to be the case here, leave the protein sufficiently functional.

**Figure 4.**
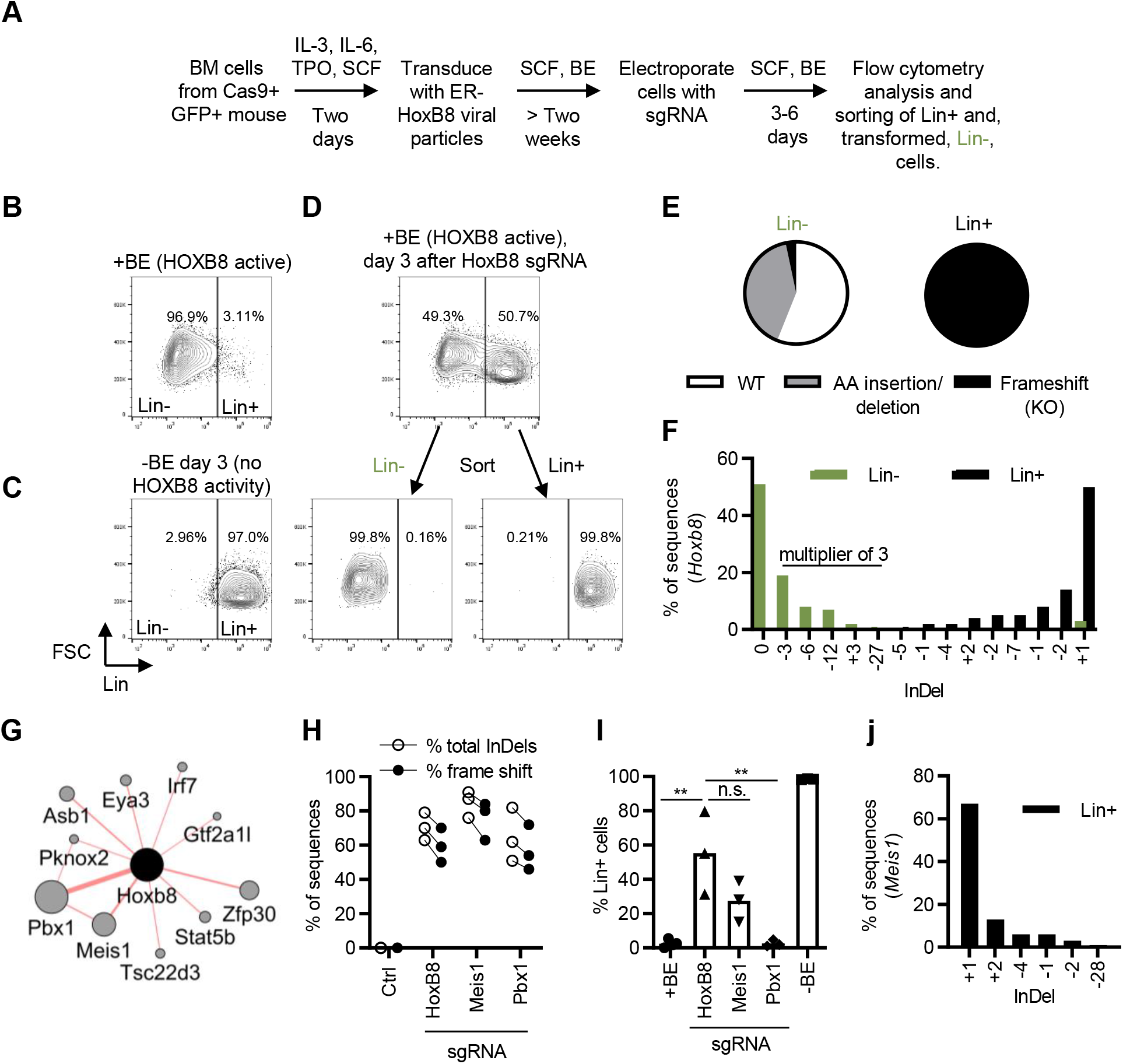
Using the Rapid CRISPR Competitive Assay (RCC) to study transformation by the HoxB8 proto-oncogene. (**A**) Model describing the experimental setup where Cas9+ BM cells were transduced with Estrogen Receptor (ER)-HoxB8 retroviral particles. HoxB8 in the transduced cells is activated by the addition of β-estradiol (BE), resulting in proliferation and block of differentiation at an immature Lin-stage. The cells were subsequently electroporated with different sgRNAs to identify genes affecting the transformed, Lin-, phenotype. (**B-C**) Flow cytometry analysis of ER-HoxB8 BM cells in the presence (B), or absence of BE (C). (**D**) Sorting of Lin- and Lin+ ER-HoxB8 BM cells three days after electroporation with a HoxB8 sgRNA. (**E**) Sequencing of the sgRNA targeted *HoxB8* locus, and deconvolution of mutation spectrum using ICE. Data shows the % of different InDels identified in the Lin-(green) and Lin+ (black) sorted cells. (**F**) representation of the type of mutations found in the sorted Lin- and Lin+ population; WT (no InDels), Amino acid (AA) insertion/deletion (InDels with a multiplier of 3 nucleotides, resulting in the addition or removal of amino acids), Frameshift (InDels with a multiplier of 1 or 2 nucleotides, resulting in a frameshift, and introduction of premature stop codon). (**G**) Top physical interaction partners of HOXB8 identified by geneMANIA. The size of the circle indicates the level of identified interactions, where the larger circles represent more prominent interaction partners. (**H-I**) The ER-HoxB8 BM cells were cultured in BE, to keep HOXB8 active, electroporated with indicated sgRNAs, and sequenced to quantify the level of induced mutations (H), as well as analyzed by flow cytometry for Lin expression (% Lin+ cells) after 3 or 6 days (three independent experiments represented in the graphs) (I). (**J**) Genotype of the CRISPR-targeted *Meis1* region in sorted Lin+ cells from Meis1 sgRNA electroporated cells. Data shown as individual data points (H, n=3), or mean and individual data points (I, n=3). n.s. = non-significant, ** = p < 0.01 by one-way ANOVA and Turkey’s post-test (I).

Since our data indicated that we could use the experimental setup to study the role of genes influencing the transformation caused by HOXB8, we set out to test whether we could identify more genes involved in the transformed phenotype. Using geneMANIA we identified a list of proteins that physically interact with HOXB8 (**Fig. 4G**). In addition to the already used Hoxb8 sgRNA, we also decided to target the top two identified candidate genes *Pbx1* and *Meis1* in HOXB8 transduced cells. We found that all three sgRNA induced a good level of mutations, with a high level of frameshift mutations in the targeted cell population (**Fig. 4H**). Moreover, we observed that the Meis1 sgRNA was able to induce the differentiation of HOXB8 transduced cells to the Lin+ phenotype, although at a lower extent when comparing to Hoxb8 targeting (**Fig. 4I**). In contrast, the Pbx1 sgRNA had no effect on the transformed Lin-phenotype despite the high level of mutations found in the cells (**Fig. 4H-I**). Sorting the Lin+ cells in the Meis1 sgRNA targeted cell population further confirmed that all identified sequences were InDels with a multiplication of one or two nucleotides, expected to generate a KO phenotype (**Fig. 4J**). We concluded that the presented experimental setup is suitable to study genes affecting the transformed state of a leukemia-like cell population and that Sanger sequencing can be used as a simple readout to evaluate the experiment.

## 5. Discussion

Comparisons of WT and KO cells have been used for many decades to dissect the role of specific genes in a given biological context. Traditionally, generating KO alleles have involved time-consuming and expensive homologous recombination techniques [49]. With the development of flexible, sequence-specific nucleases, including CRISPR, this process has been dramatically simplified [50]. However, the resulting genotype in a CRISPR-targeted cell population is not uniform, as exemplified in Fig. 1F. With a binary mindset (e.g. WT or KO), the analysis of a CRISPR-targeted population could thus be non-productive. Often, researchers address the genetic heterogeneity by generating clonal lines of the targeted cells or animals. This is however not always possible, can result in the selection of traits that are not directly linked to the intended genotype, and is also time consuming. Instead of identifying the CRISPR-induced genetic heterogeneity as a problem, we here hypothesized that the heterogeneity could be embraced for discovery, and that regular Sanger sequencing could be used as a simple readout to quantify the heterogeneity and identify genotype enrichment. For this purpose, genomic DNA was isolated from the cells, the CRISPR-targeted region amplified by PCR and sequenced by standard Sanger sequencing approach followed by analyzing the sequencing result file with the free web-based software ICE [13]. Delivering a GFP targeting sgRNA into the Lin-BM cells, isolated from Cas9+ GFP+ mice, we compared a flow cytometry-based assay for GFP inactivation to the sequencing of the targeted GFP locus. We found that the readouts showed a good correlation (R^2^=0.87), although the sequencing readout evidenced a lower sensitivity when the mutations were found at a low frequency (Fig. 1G). This could be expected based on the way ICE analyzes the samples, where the mixed peaks of the sequencing readout are deconvoluted into frequencies, with low-frequency mutations thus being more difficult to resolve. As an alternative readout, we used the same genomic DNA samples and the same primers to perform fragment length analysis (FLA) using the IDAA technology [11, 12]. This approach separates the PCR product by capillary electrophoresis and in a precise way defines the size of the different products formed in the PCR reaction. This approach is not constrained by the same type of sensitivity issues as the sequence deconvolution approach and showed a great correlation (R^2^=0.99) to the flow cytometry-based readout (Fig. 1H). Based on the simplicity, speed, and cost-effectiveness of the Sanger sequencing approach, we have continued to use this readout, keeping in mind the limitation of detection at low mutation frequencies. Notably, since the same genomic DNA samples and primers can be used for the FLA, important samples can be analyzed first by the Sanger sequencing approach and subsequently by FLA if necessary. NGS-based readouts can also be considered, and several amplicon-based analysis pipelines are established for analyzing CRISPR targets [8-10].

As we had optimized culture conditions and sgRNA delivery to mouse Lin-BM cells, we next performed a set of experiments to assess the discovery potential using Sanger sequencing as a readout. Conceptually, the idea was to let cells with different genotypes compete and see if specific genotypes were enriched when studying different cellular behaviors or phenotypes. We refer to this setup as the rapid CRISPR competitive (RCC) assay. Notably, in many ways, the RCC assay exploits the same fundamental mechanisms as a CRISPR screen but focuses on one gene (that is Sanger sequenced), instead of a set of genes (where sgRNA barcodes are sequenced by NGS in the screen setting). To this end, we first tested if the approach could be used to identify genes that affect the development of immune cells in vivo. We electroporated the Lin-BM cells (GFP+ Cas9+ CD45.1+) with a Zap70 sgRNA and transfer them into irradiated CD45.2+ (GFP+ Cas9+ CD45.2+) recipient mice. In the recipient mice, the transferred, modified BM cells graft into the BM compartment and start generating new immune cells that can be tracked by the CD45.1 expression. We refer to these mice as immuno-CRISPR (iCR) mice. *Zap70* was selected as a proof-of-concept target, as it is known to be essential for T cell development, while not affecting for example B cell development [39-41]. As anticipated, we found that sorted CD45.1+ B cells had a high proportion of mutations in *Zap70*, while we could not detect any mutations in CD45.1+ T cells in the Zap70 iCR mice (Fig. 2D). The RCC assay thus worked well to evaluate the role of *Zap70 in vivo*. Considering the complexity and cost of generating gene-modified mice, even with novel CRISPR-based approaches, we see great potential in using the iCR approach to rapidly study the role of different candidate genes in mature immune cells and hematopoiesis. This approach shares similarities with mixed bone marrow chimera experiments, but instead of using flow cytometry to identify the enrichment/depletion of cells with specific congenic markers (used as a proxy for a specific genotype), sequencing is directly used to identify the enrichment/depletion of specific genotypes.

In line with the *in vivo* differentiation data, we also found that the modified Lin-BM cells could be readily differentiated *in vitro* into both macrophages and dendritic cells with an expected phenotype (Fig. 3B, F). Importantly, we found that in a mixed population of GFP WT and GFP KO macrophages differentiated from BM cells electroporated with a GFP sgRNA, both cells performed equally well in a functional phagocytosis assay (Fig. 3D-E). As GFP is not involved in the phagocytosis process, this experiment shows that CRISPR-induced DNA damage does not compromise or generate non-specific effects in the target cells [34-36].

Lastly, we explored whether our analysis pipeline could also be used to study malignant transformation induced by the overexpression of a proto-oncogene. We induced the activity of HOXB8 in BM cells, resulting in a transformed state characterized by the cells proliferating at an immature (Lin-), leukemia-like, differentiation stage [29]. Subsequently, we evaluated cell differentiation into the mature (Lin+) non-transformed state as cells were electroporated with different sgRNAs, aiming to identify potential drug targets affecting the transformed phenotype. Initially, we targeted *Hoxb8* itself as a proof-of-concept. We found that half of the CRISPR-targeted cells lost the leukemia-like phenotype and differentiated to mature (Lin+) cells defined by expected inactivating mutations in the *Hoxb8* region (Fig. 4D-F). Interestingly, we also found that the Lin-population had a fair amount of mutations, despite retaining the HOXB8-induced transformed phenotype (Fig. 4E-F). By closer examination, we noted that the mutations found in the Lin-population were mainly -3, -6, and-12 nucleotides, corresponding to the deletion of 1, 2, and 4 AAs, respectively. In the Lin+ population, we instead found a dominant +1 mutation, followed by less abundant -2, -1, -7, -2, +2, -4 and -1 mutations, all being frameshift mutations resulting in premature stop codons, and nonsense-mediated decay [47, 48]. This observation suggests that the deleted AAs found in the Lin-population are not essential for the HOXB8 activity. Arguably, such a phenomenon can be expected to be very protein and target specific. For example, we see no evidence for such phenomenon with the used GFP sgRNA, where sorted GFP+ cells in a population targeted by the GFP sgRNA, had no detectable InDels (Fig. 1J). Nevertheless, the Hoxb8 data (Fig. 4E-F) identified that comparing the frequency of total InDels to the frequency of InDels expected to result in a KO phenotype (insertion/deletion with a multiplier of 1-2 nucleotides; frameshift) could be a way to identify protein domains with structural and functional significance.

The lack of effect by knocking out *Pbx1* in HOXB8 overexpressing cells (Fig. 4I) was surprising as the PBX1 binding site in HOXB8 has been reported to be important for most, but not all, features of HOXB8 overexpression in experimental systems [46]. However, our data is in line with CRISPR screen data from HOXB8 overexpressing cells, identifying that *Hoxb8*, and *Meis1*, but not *Pbx1*, sgRNAs are lost over time [51]. The influence of *Pbx1* deficiency on the system could also be influenced by the specific differentiation stage of the HOXB8 transformed cells, where the SCF culture condition used here results in a granulocyte-macrophage progenitor (GMP) phenotype. Notably, Ficara et al. showed that *Pbx1* deficiency gives a relatively mild phenotype in GMP cells compared to for example long-term hematopoietic stem cells (LT-HSCs) [52].

The concept of comparing cells or microorganisms with different genotypes in competitive settings is a proven discovery model. It is, for example, the basis for CRISPR and shRNA screens, as well as for mixed BM chimera experiments. The same “survival of the fittest” mechanisms is furthermore the basis for the enrichment of specific mutations in cancer cells and infectious agents, both spontaneously over time and in response to drugs [53-56]. The selection of specific genotypes in all these settings infers a central functionality to the specific genotypes and can thereby guide the development of drug candidates targeting the identified genes/proteins. Here, we set out to establish a simple experimental setup to identify the role of different genes in studied cellular behaviors. We use standard Sanger sequencing to quantify the genetic diversity induced at a CRISPR-targeted site and use enrichment of specific genotypes to identify the role for the studied gene.

## 6. Conclusions

Sanger sequencing and sequence deconvolution can be used as a rapid discovery readout to identify the role of CRISPR -targeted genes.

## Abbreviations

(AA): Amino acid
(BM): bone marrow
(CRISPR): Clustered Regularly Interspaced Short Palindromic Repeats
(ctrl): control
(FLA): fragment length analysis
(GMP): granulocyte-macrophage precursor
(HSC): hematopoietic stem cell
(iCR): immuno-CRISPR
(IDAA): Indel Detection by Amplicon Analysis
(ICE): Inference of CRISPR Edits
(InDel): insertion or deletion
(KO): knockout
(Lin): lineage
(NGS): next-generation sequencing
(PBMC): peripheral blood mononuclear cells
(RCC assay): rapid CRISPR competitive assay
(sgRNA): single guide RNA
(shRNA): short hairpin RNA
(siRNA): small interfering RNA
(TIDE): Tracking of Indels by Decomposition
(TRAC): T cell receptor alpha chain constant
(WT): wild type

## 7. Acknowledgements

We are grateful to Drs. Petter Woll, Helena Malmgren, Lisa Westerberg, Taras Kreslavskiy, and Sudeepta Panda for valuable discussions. The ER-Hoxb8 construct was a gift from Mark P. Kamps, University of California, San Diego. The pSIRV-NFAT-eGFP construct was a gift from Peter Steinberger, Medical University of Vienna. This research was partly funded by grants from the Swedish Research Council, the Swedish Cancer Society, Karolinska Institutet, Åke Olssons stiftelse, Magnus Bergvalls stiftelse, Stiftelsen Professor Nanna Svartz fond, Felix Mindus contribution to Leukemia Research (to FW), the China Scholarship Council (to LJ and YS), and the Nanyang Technological University–Karolinska Institutet Joint PhD Programme (to VSI).

---

**Supplementary Figure 1.**
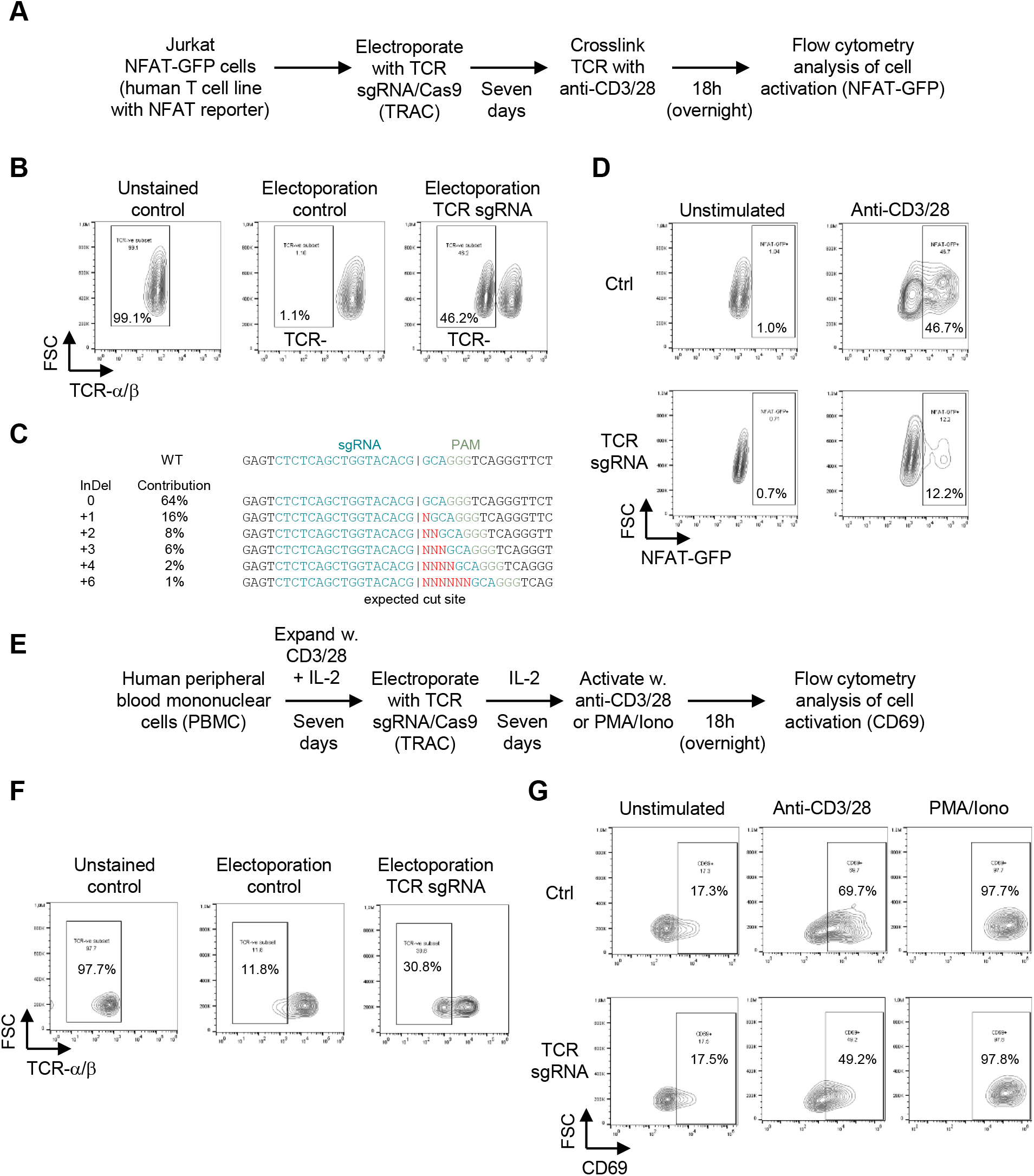
CRISPR KO in T cells. (**A**). Model describing the experimental setup where Jurkat NFAT-GFP reporter cells were electroporated with a sgRNA/Cas9 complex targeting the constant domain of the TCR alpha chain (TRAC). (**B-C**) Confirmation of TCR KO efficiency in the Jurkat NFAT-GFP cells by flow cytometry (anti-TCR-α/β-APC) (B), and by sequencing (C) two days after electroporation. Unstained control was not electroporated. Electroporated control was electroporated without a sgRNA. (**D**) TCR sgRNA treated cells show a dampened response to TCR stimulation through lesser GFP upregulation following activation. CD3/28 stimulation was done with CD3/28 coated magnetic beads for 24 hours in a 1:1 ratio to the total number of cells. (**E**) Model describing the experimental setup where human peripheral blood mononuclear cells (PBMC) were electroporated with the TRAC sgRNA/Cas9 complex. (**F**) Confirmation of the level of TCR KO in PBMCs by flow cytometry (anti-TCR-α/β-APC). Unstained control was not electroporated. Electroporated control was electroporated without a sgRNA. (**G**) TCR sgRNA treated PBMC shows a dampened response to TCR stimulation through lower CD69 upregulation. As expected, the TCR sgRNA does not affect alternative activation pathways as in the case of PMA/Ionomycin stimulation. CD3/28 stimulation was done with CD3/28 coated magnetic beads for 18 hours in a 1:1 ratio to the total number of cells. PMA/Ionomycin stimulation was done with 100nM/mL each of PMA and Ionomycin for 18 hours.

**Supp. Table 1.**
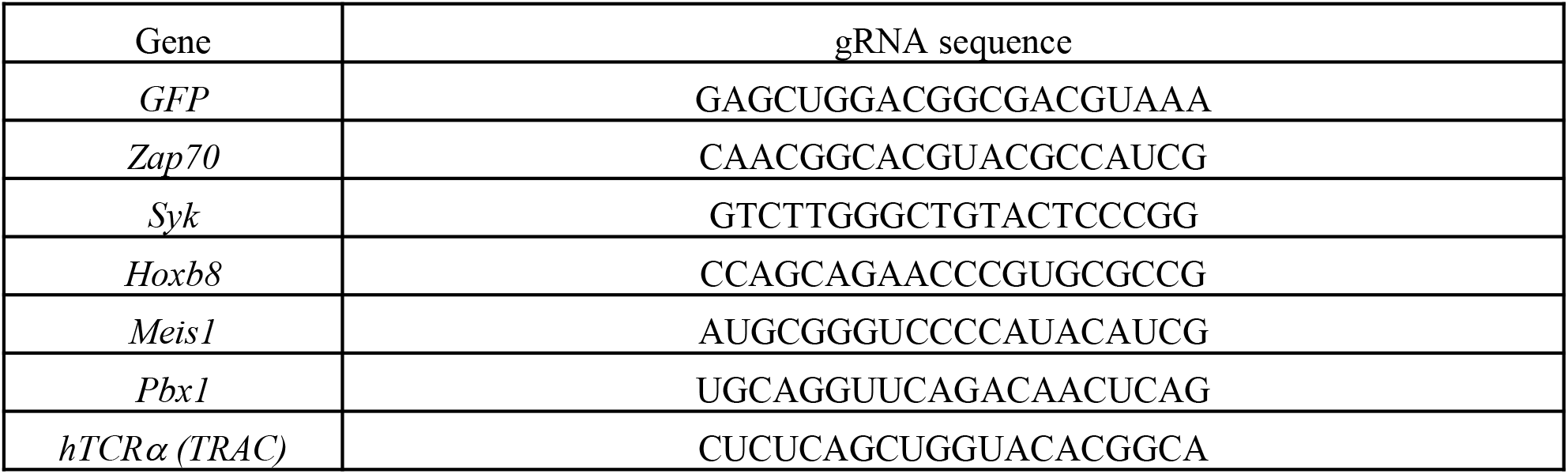

**Supp. Table 2.**
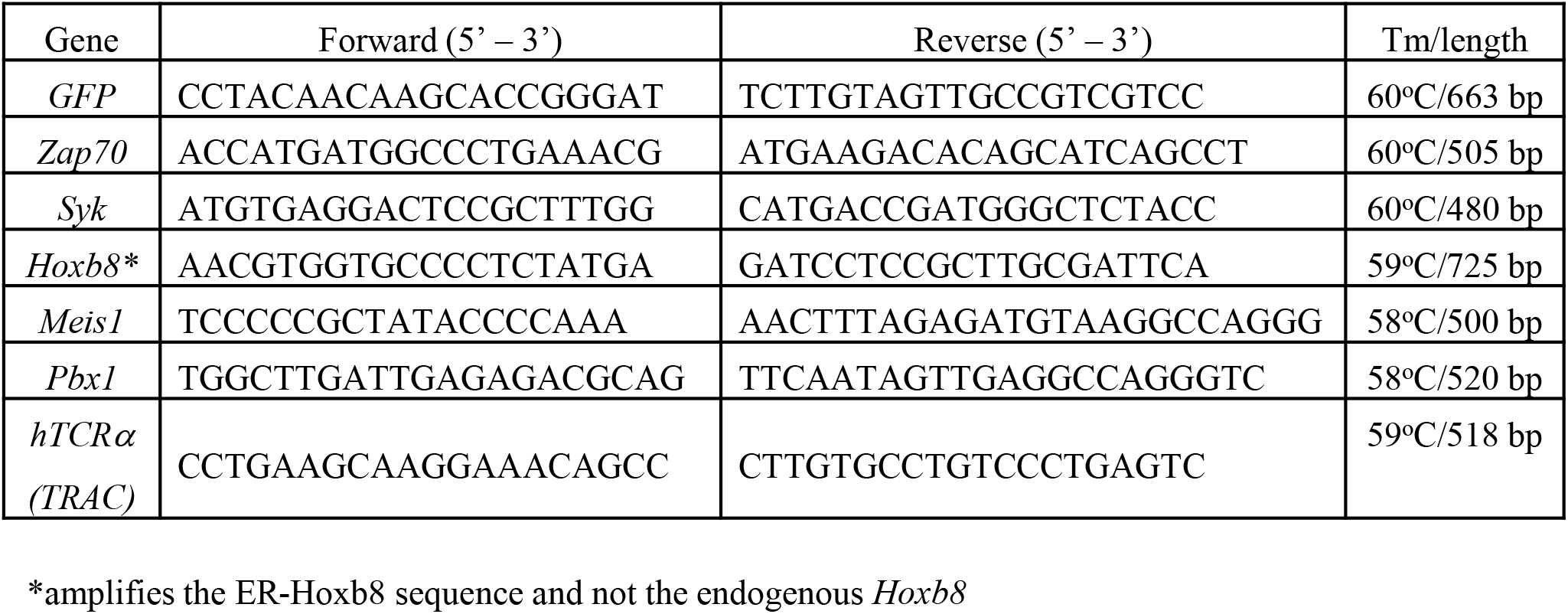

